# Integration of Nuclear Receptors into a Schwann cell Gene Regulatory Network

**DOI:** 10.1101/2025.10.30.685409

**Authors:** Raghu Ramesh, Saniya Khullar, Alena J. Hanson, Seongsik Won, Camila Lopez-Anido, Daifeng Wang, John Svaren

## Abstract

Peripheral nerve function depends on the proper formation and maintenance of Schwann cells. During development, large diameter axons are enveloped by single, myelinating Schwann cells whereas clusters of small diameter axons are surrounded by nonmyelinating Schwann cells in Remak bundles. The transcription factor SRY-box 10 (SOX10) is required for the development of Schwann cells, but its functions in different neural crest and glial lineages have shown that SOX10 commonly cooperates with other transcription factors to activate cell-specific gene programs.

In a previous study, we compared SOX10-bound regulatory elements in Schwann cells and identified a specific enrichment for nuclear receptor motifs at SOX10 binding sites in Schwann cells. NR2F1 and NR2F2 (Coup-TFI/Coup-TFII) are expressed from neural crest through Schwann cell maturity, and knockdown of nuclear receptors *Nr2f1* and *Nr2f2* in primary Schwann cells downregulated several myelin genes. In this study, we have identified a NR2F-regulated target gene network in Schwann cells, which revealed enrichment for nonmyelinating Schwann cell genes. Cut&Run assays in S16 Schwann cells revealed novel, genome-wide binding sites of NR2F2 and two downstream transcription factors: 1) Retinoid X Receptor Gamma (RXRG), which is also expressed preferentially in nonmyelinating Schwann cells, and 2) TEA-Domain factor (TEAD1), which is an important HIPPO pathway component required for Remak bundle formation. Our study elucidates the transcriptional cooperation that programs the regulatory network in nonmyelinating Schwann cells.

## Introduction

Schwann cells enable saltatory conduction in peripheral nerve and are required for axonal homeostasis (Poitelon et al., 2020; Taveggia and Feltri, 2022). Myelin produced by Schwann cells envelops segments of single, large-diameter axons to drastically reduce membrane capacitance, accelerate nerve conduction, and facilitate appropriate sensory and motor responses (Quintes et al., 2010; Poitelon et al., 2020). In contrast, nonmyelinating Schwann cells (nmSCs) wrap several small diameter axons to form Remak bundles (Huang et al., 2005; Griffin and Thompson, 2008; Harty and Monk, 2017). Axoglial signaling is important for both types of Schwann cells (Nave and Salzer, 2006; Taveggia et al., 2010), but nonmyelinating Schwann cells play critical roles in modulation of several types of processes, including initiation and modulation of nociception (Poplawski et al., 2018; Abdo et al., 2019; Rinwa et al., 2021; Martellucci et al., 2025). For example, nonmyelinating Schwann cells regulate sensory neuron activity in Fabry disease (Waltz et al., 2024), produce prostaglandin E2 to modulate sensory axons (Kantarci et al., 2024), and even provide an important role in hematopoietic stem cell maintenance in bone marrow (Yamazaki et al., 2011).

Single-cell studies have revealed distinct transcriptional profiles of myelinating vs. nonmyelinating Schwann cells (mSC vs. nmSC) (Gerber et al., 2021; Avraham et al., 2022; Yim et al., 2022), but much more is known regarding transcriptional regulation of embryonic Schwann development and formation of myelinating cells(Parkinson and Svaren, 2020), with relatively little information on coordination of gene regulatory networks in nonmyelinating cells. A recently generated computational framework based on human single cell data has provided some insight into the distinct transcriptional networks of myelinating vs. nonmyelinating Schwann cells (Khullar et al., 2025).

A key transcription factor (TF) involved in Schwann cell maturation is the SRY-box10 (SOX10) protein, which is expressed at several stages of Schwann cell development and is required for myelin formation and maintenance (Finzsch et al., 2010; Bremer et al., 2011; Sock and Wegner, 2019). In humans, mutations of the *SOX10* gene cause a neurocristopathy syndrome known as PCWH: peripheral demyelinating neuropathy, central dysmyelinating leukodystrophy, Waardenburg syndrome, and Hirschsprung disease (Verheij et al., 2006). SOX10 associates with other transcription factors important for Schwann cell development such as the paired domain transcription factor PAX3, POU domain protein POU3F1/OCT6/SCIP, and early growth response factor EGR2 (Reiprich et al., 2010; Srinivasan et al., 2012; Wahlbuhl et al., 2012). Mutations in the *EGR2* gene cause a severe neuropathy in humans, while *Egr2 (Krox20)* -/- mice exhibit a dramatic lack of myelin formation (Topilko et al., 1994; Warner et al., 1998; Warner et al., 1999; Le et al., 2005; Lupo et al., 2020).

SOX10 is expressed in several other cell types such as oligodendrocytes (Li et al., 2007; Liu et al., 2007), but relatively few SOX10 binding sites are shared in Schwann cells vs. oligodendrocytes, suggesting that SOX10-regulated enhancers arise in a cell specific manner using unique combinations of transcription factors (Bujalka et al., 2013; Lopez-Anido et al., 2015). Indeed, neither SOX10 nor any other TF is exclusively expressed in Schwann cells vs. other cell types, therefore our goal is to identify the unique combinations of transcription factors that drive Schwann cell differentiation in accord with the well-established combinatorial regulation of transcription (Arnone and Davidson, 1997).

Our comparative analysis of SOX10-bound enhancers in Schwann cells vs. oligodendrocytes identified NR2F nuclear receptors (NR2F1/COUP-TFI and NR2F2/COUP-TFII) as candidate regulators of Schwann cell differentiation (Lopez-Anido et al., 2015). NR2F1 and NR2F2 are expressed from neural crest to mature Schwann cells (Rada-Iglesias et al., 2012; Kastriti et al., 2022). An analysis of human stem cell-derived neural crest revealed a co-enrichment of NR2F1 and NR2F2 binding sites along with other neural crest transcription factors such as SOX10 and TFAP2A, and loss-of-function experiments in human and Xenopus neural crest cells indicated an important role in neural crest development (Rada-Iglesias et al., 2012). We found that siRNA-mediated reduction of *Nr2f1* and *Nr2f2* in Schwann cells reduced transcript levels for *Mbp*, *Ndrg1*, and *Dhh*, which are all expressed by Schwann cells in peripheral nerve (Martini et al., 1995; Parmantier et al., 1999; Heller et al., 2014).

Nuclear receptors NR2F1 (COUP-TFI) and NR2F2 (COUP-TFII) play important roles in nervous system development (Yamaguchi et al., 2004; Rada-Iglesias et al., 2012; Olivares et al., 2015; Polvani et al., 2019). Although there are no reported studies of NR2F mutants affecting peripheral nerve, loss of function mutations in *NR2F1* lead to Bosch-Boonstra-Schaff Optic Atrophy Syndrome (BBSOAS) which is characterized by vision loss, impairment of intellectual development, and brain malformations leading to autism and seizures (Tocco et al., 2021). *Nr2f1* null mice exhibit delays in oligodendrocyte differentiation, and demyelination in the central nervous system (Yamaguchi et al., 2004; Bertacchi et al., 2019). Heterozygous knockouts of *Nr2f1* in mice show characteristics of BBSOAS (Chen et al., 2020), including imbalanced populations of astrocytes versus oligodendrocytes leading to hypomyelination of the optic nerve, among other defects (Bertacchi et al., 2019).

Since SOX10 interacts with binding partners to promote myelinating cell fates (Kuhlbrodt et al., 1998a; Schreiner et al., 2007; Stanchina et al., 2010), we have explored how NR2F family members may interact with SOX10 and other transcription factors to co-regulate Schwann cell genes.

## Results

### NR2F1 and NR2F2 Regulate Nonmyelinating Schwann Cell Genes

Since our preliminary study in primary rat Schwann cells indicated that NR2F1 and NR2F2 regulate some Schwann cell genes involved with development and myelination (Lopez-Anido et al., 2015), we tested for interactions of NR2F1 and NR2F2 with SOX10 and other transcription factors using the S16 Schwann cell line. This cell line expresses high levels of myelin genes (Sasagasako et al., 1996; Hai et al., 2002), and many of the enhancers that have been identified in peripheral nerve *in vivo* are present in S16 cells (Jang and Svaren, 2009; Jones et al., 2012; Srinivasan et al., 2012). However, they also express several nmSC cell genes (e.g. *Ngfr* and *Gfap*) in a SOX10-dependent manner (Fogarty et al., 2020). We treated S16 Schwann cells with siRNAs for *Nr2f1* and *Nr2f2* together, and performed RNA-seq to identify candidate downstream gene targets (Supporting Information Table 1 and Figure S1).

**Table 1.**
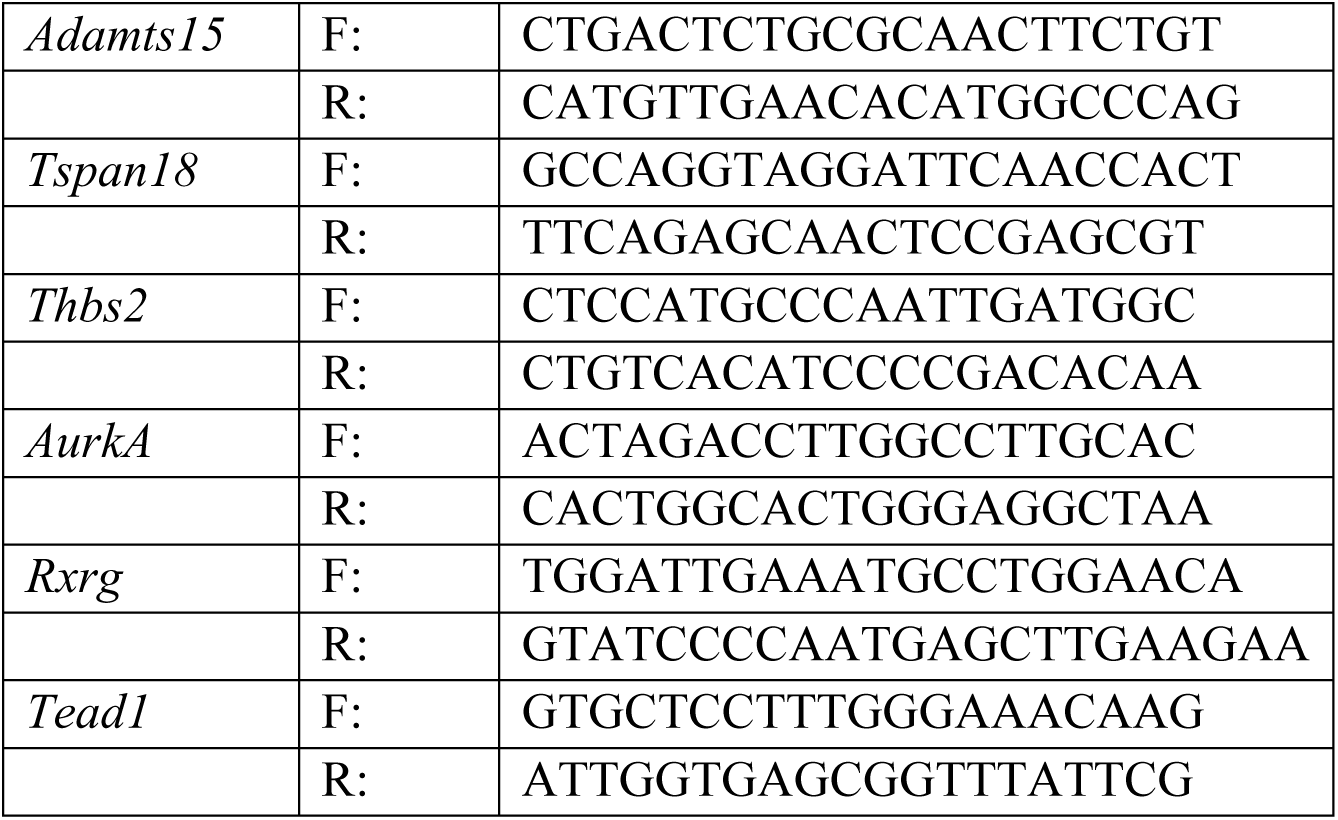
Primer Sequences Used for qRT-PCR Experiments.

The results confirmed downregulation of *Nr2f1* and *Nr2f2*, and revealed modulation of a major subset of genes expressed in Schwann cells, with upregulation of genes such as fatty acid 2 hydroxylase (*Fa2h*) and significant downregulation of genes such as *Adamts15*, as shown in the volcano plot (Fig. 1A). Since NR2F1 and NR2F2 factors can be gene activators, we first took a subset of the most significantly downregulated genes in the presence of *Nr2f1* and *Nr2f2* siRNA and verified that many of the downregulated genes are expressed in Schwann cells *in vivo* by comparison to recently published analyses of single-cell RNA-seq and sorted Schwann cells (Gerber et al., 2021; Brosius Lutz et al., 2022; Yim et al., 2022). Gene ontology analyses of downregulated genes revealed cell cycle, focal adhesion and HIPPO signaling pathways (Supporting Information Figure S2B).

**Figure 1.**
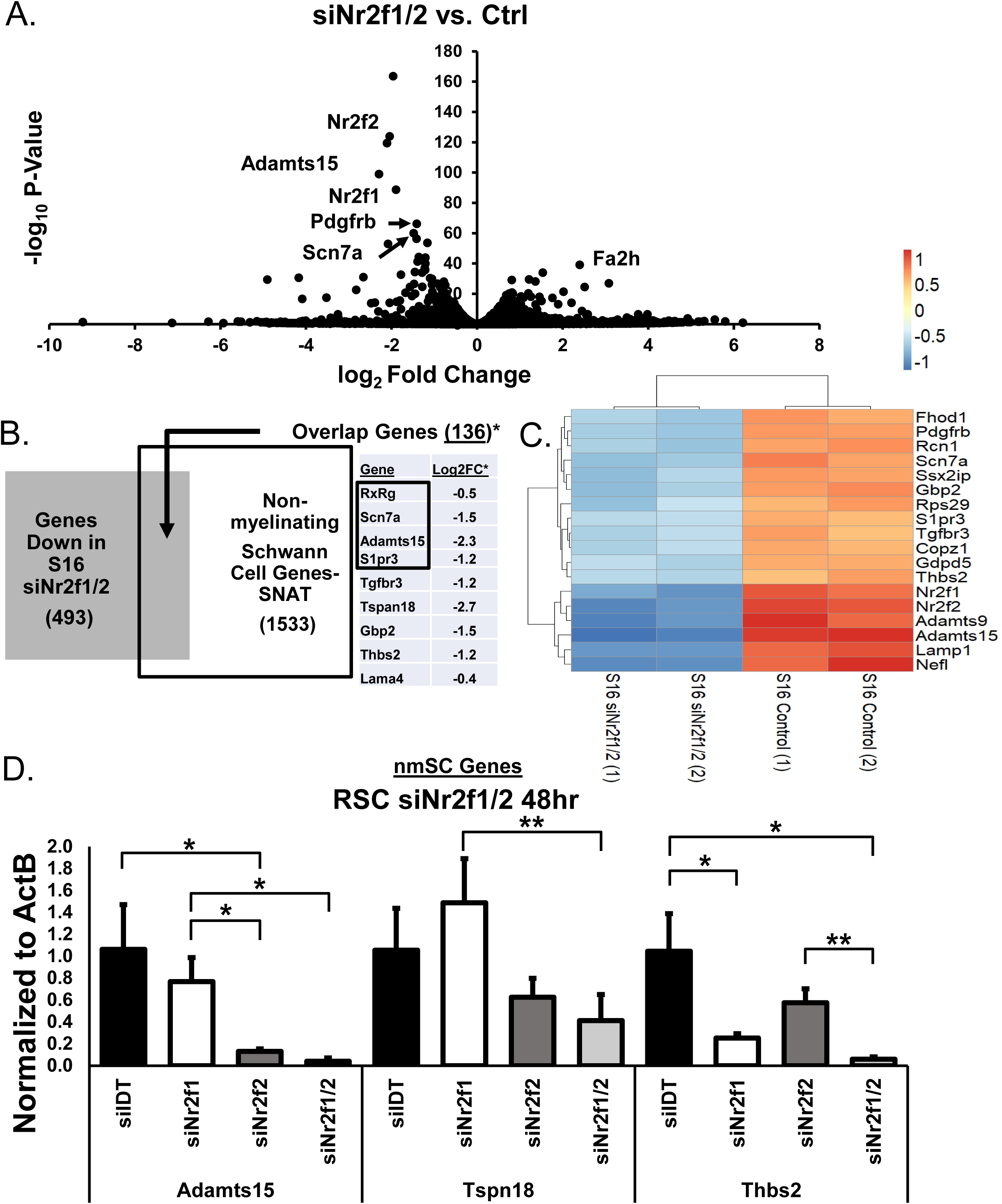
Downstream Gene Targets of NR2F1/2 Regulation. **(A)** Volcano plot from RNA-seq data of S16 Schwann cells treated with siRNA for *Nr2f1* and *Nr2f2* relative to control siRNA (48 hr). Most strongly changed genes are downregulated, including *Nr2f1* and *Nr2f2* transcripts. **(B)** Overlap analysis between genes downregulated in the *siNr2f1/2* RNA-seq data (Adj. P-Value < .05, ≤.5 log2FC) versus nonmyelinating Schwann cell genes preferentially expressed in the Sciatic Nerve Atlas [nmSC (Avg. > RPKM 20) / mSC)) = DE (>2 fold)]. Table lists the top 9 most significantly downregulated genes from the *siNr2f1/2* RNA-seq data that overlap with differentially expressed nonmyelinating Schwann cell genes from the Sciatic Nerve Atlas (* = p < 0.05). Black box highlights the most differentially expressed nonmyelinating Schwann cell genes over myelinating Schwann cell genes from the Sciatic Nerve Atlas. **(C)** RPKM values from the top 20 downregulated genes by adjusted p-value in the siNr2f1/2 RNA-seq data were log2 transformed, sorted by row means, and clustered based on differential expression trends between control and knockdown conditions. **(D)** qRT-PCR analysis of primary rat Schwann cells treated with *Nr2f1/2* siRNA for 48 hr (n=3) shows downregulation of a subset of nonmyelinating Schwann cell genes.

Since NR2F1 and NR2F2 were initially identified in an analysis of SOX10 target genes (Lopez-Anido et al., 2015), we accordingly found a significant overlap of 115 NR2F1 and NR2F2-regulated genes out of the 1390 SOX10-regulated genes that had been identified in a previous study of a S16 Schwann cell SOX10 CRISPR knockout (Fogarty et al., 2020) (Supporting Information Table S3). In the course of the analysis, we found that relatively few of the NR2F1 and NR2F2-regulated genes are expressed in mSC. Instead, some of the most downregulated genes are preferentially expressed in nmSC (Deerinck et al., 1997; Watanabe et al., 2002; Gerber et al., 2021; Yim et al., 2022).

To define these genes further, we compared the significantly downregulated genes in our *siNr2f1/2* RNA-seq data to genes enriched in nmSC based on recent gene expression profiling of myelinating vs. nonmyelinating Schwann cells (Gerber et al., 2021). Out of the 1533 nmSC enriched genes from the Sciatic Nerve Atlas, 136 overlap with the 493 significantly downregulated genes in our RNA-seq data (p < 0.01, Fig. 1B and Supporting Information Table S4). A heatmap of the *siNr2f1/2* RNA-seq data clusters shows *Adamts15* and *Tspan18* as genes that are strongly and similarly downregulated upon reduction of *Nr2f1* and *Nr2f2* (Fig. 1C). Selected nmSC genes were also tested in primary rat Schwann cells, and we found several genes that were similarly regulated by *Nr2f1* and *Nr2f2* (Fig. 1D). To address the possibility of off-target effects, we used an independent pair of Nr2f1 and Nr2f2 siRNA’s in the S16 cell line and saw a similar and overlapping reduction of nmSC genes (Supporting Information Table S2 and Figure S3).

### NR2F2 Colocalizes with SOX10 at Nonmyelinating Schwann Cell Gene Enhancers

The nuclear receptor motifs are enriched in SOX10-bound enhancers in Schwann cells compared to oligodendrocytes (Lopez-Anido et al., 2015). Therefore, we tested if NR2F1/2 proteins localize to SOX10-bound enhancers using Cut&Run assays (Skene and Henikoff, 2017) for NR2F2 in S16 Schwann cells. In addition, we performed Cut&Run analysis for the active enhancer mark, H3K27 acetylation (Rada-Iglesias et al., 2011), to identify active enhancers in the S16 Schwann cell line.

We compared our S16 Schwann cell NR2F2 peak data against SOX10-bound active enhancers in vivo (Lopez-Anido et al., 2015). The heatmaps show NR2F2 read density over the SOX10-bound enhancers defined previously (Fig. 2A). Overall, NR2F2 binds to 15.3% of the SOX10-bound Schwann cell enhancers in the S16 cell line. This overlap suggests that many nmSC genes are regulated by cooperation of NR2F1 and NR2F2 with SOX10. In contrast, we compared active S16 enhancers that overlap with oligodendrocyte SOX10-bound active enhancers (Lopez-Anido et al., 2015), but only 5.9% of these enhancers interact with NR2F2. This is consistent with previously described enrichment of nuclear receptor motifs in Schwann cell vs. oligodendrocyte enhancers (Lopez-Anido et al., 2015). We also analyzed binding of NR2F1 (not shown), but it is expressed at lower levels than NR2F2 in Schwann cells (Fogarty et al., 2020) and the read density pattern over SOX10-bound enhancers was less distinct compared to NR2F2, although it showed binding to similar sites.

**Figure 2.**
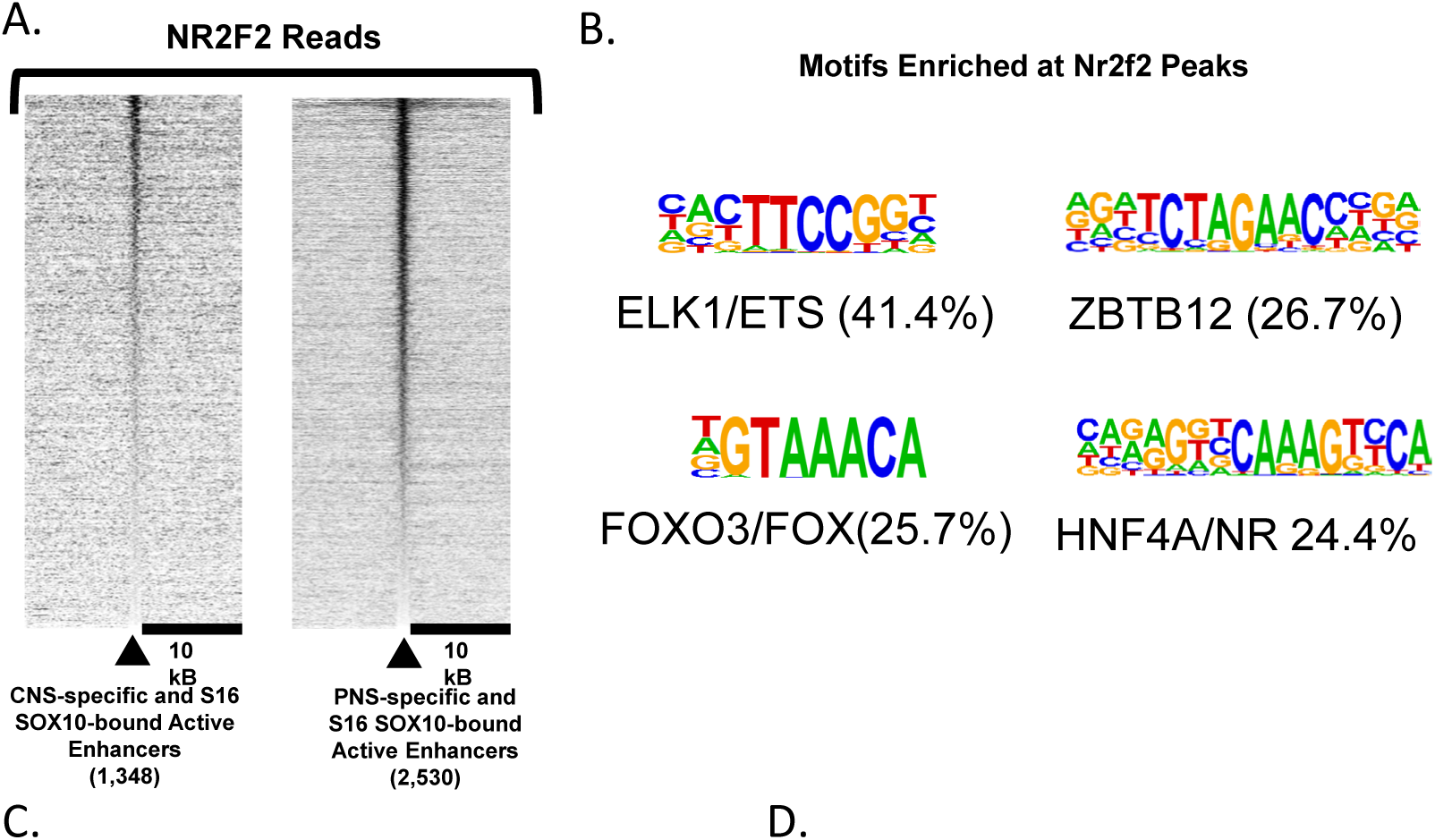
Cut&Run Assays of NR2F2 Binding. **(A)** Heatmaps of NR2F2 read density in S16 Schwann cells centered at peaks of SOX10-bound active enhancers in the central nervous system (spinal cord) and peripheral nervous system (sciatic nerve) show that NR2F2 reads are preferentially distributed around PNS SOX10-bound active enhancers. **(B)** HOMER motif analysis of NR2F2-bound enhancers reveals strong enrichment for ETS, FOX, zinc-finger (ZF), and the nuclear receptor motif bound by NR2F-related family members (NR, HNF4A)**. (C)** qRT-PCR analysis of S16 Schwann cells treated with siRNA for *Nr2f1*, *Nr2f2*, and *Sox10* for 48 hours shows combinatorial regulation of three nmSC genes by the triple knockdown (*Nr2f1/Nr2f2/Sox10*). **(D)** S16 Schwann cell Cut&Run tracks for NR2F2 and the active enhancer mark H3K27ac, coupled with the sciatic nerve ChIP-seq track for SOX10, show colocalization of SOX10 and NR2F2 at active nonmyelinating Schwann cells enhancers for genes *Ncam1*, *Nav2*, and *S1pr3*. Red boxes show called peaks of significant binding.

A motif analysis (Heinz et al., 2010) was performed on the called peaks from NR2F2 Cut&Run binding data to screen for other potential binding partners (Fig. 2B). The nuclear receptor motif (HNF4A, another NR2F family member) was among the most highly enriched across NR2F2 binding sites, but there was also enrichment of ETS and FOX family motifs. Notably, ETS and FOX family members *Ets1*, *Etv5*, *Etv6*, and *Foxp1*, *Foxo1* are preferentially expressed at the transcript level in nmSC (Gerber et al., 2021). Moreover, NR2F2 has been shown to bind to FOXA1 (Wang et al., 2020b) and FOXP1/2 (Estruch et al., 2018), and an analysis of endothelial enhancers showed a similar colocalization of NR2F2 with ETS and FOX motifs (Sissaoui et al., 2020). To determine if some target genes are jointly regulated by *Nr2f1/Nr2f2/Sox10*, we used siRNA to knockdown *Nr2f1/Nr2f2* together, *Sox10* alone, and all three together (*Nr2f1/Nr2f2/Sox10).* We considered the transcription factors to be working together when the triple knockdown (*Nr2f1/Nr2f2/Sox10*) had a significantly lower log2 fold change (p<0.05) than *siSox10* and *siNr2f1/2* individually. We show three such genes that are regulated by the combination of *Nr2f1*, *Nr2f2*, and *Sox10*: Ncam1, Nav2, and S1pr3 (Fig. 2C). All three are preferentially expressed in nmSCs (Gerber et al., 2021; Yim et al., 2022), and we identified overlapping binding sites for SOX10, NR2F2, and H3K27ac (marker of active enhancers) (Fig. 2D) around these genes.

To determine if NR2F2 is expressed in nmSC *in vivo*, we performed immunohistochemistry on cross sections of adult rat sciatic nerve for NR2F2 along with the NCAM1 cytoplasmic marker of nmSC. We found staining for NR2F2 present at nmSC nuclei (Fig. 3A-B), consistent with the preferential expression of *Nr2f2* in nmSC (Gerber et al., 2021; Yim et al., 2022).

**Figure 3.**
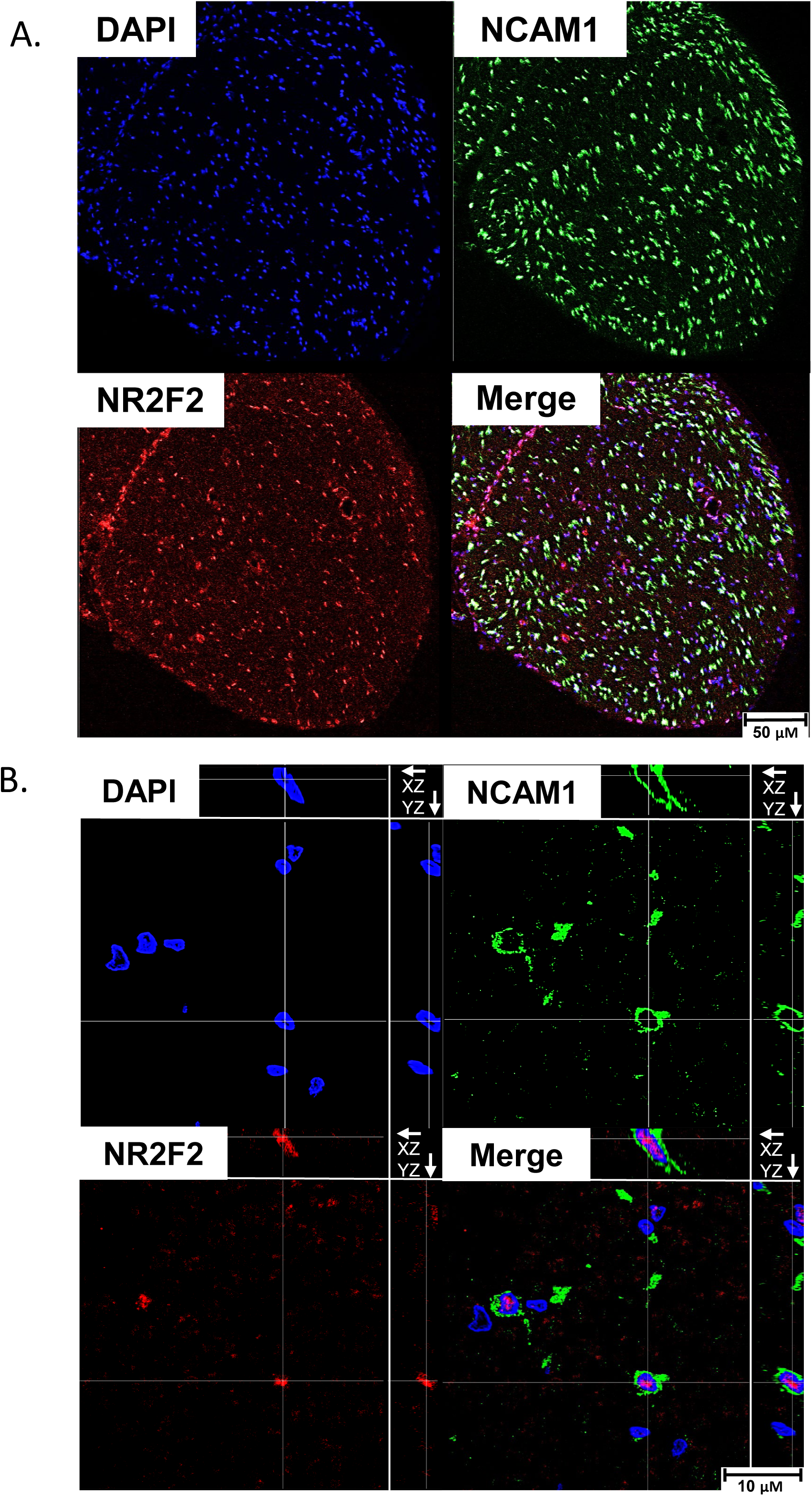
NR2F2 Localizes at Nonmyelinating Schwann Cell Nuclei *in vivo*. **(A)** Immunohistochemistry of rat sciatic nerve cross-sections at 20x using antibodies for NR2F2 (red), and a nonmyelinating Schwann cell marker NCAM1 (green), and DAPI (blue). **(B)** 63x images with orthogonal cross-sections along the x-z and y-z axis show co-localization of NR2F2 throughout distinct nonmyelinating Schwann cell nuclei.

### TEAD1 is a Downstream Target and Colocalizing Factor of NR2F2

In our previous analysis of active Schwann cell enhancers (Hung et al., 2015; Lopez-Anido et al., 2015), we had also found enrichment for Ets-domain factors (ETS), and TEA-domain transcription factors (TEAD) binding motifs. TEAD1 is a DNA-binding partner of coactivators YAP and TAZ as part of the HIPPO pathway, which is required for development of nmSC through axonal sorting and initiation of Schwann cell myelination (Poitelon et al., 2016; Grove et al., 2017; Grove et al., 2024). We also observed significant downregulation of *Tead1* in our *Nr2f1* and *Nr2f2* siRNA experiment in S16 cells (Fig. 1). Treatment of primary rat Schwann cells with siRNA for *Nr2f1* and *Nr2f2* also reduced levels of the *Tead1* transcript (Fig. 4A). Additionally, we observed binding of SOX10 and NR2F2 at an intragenic enhancer within the *Tead1* gene (Fig. 4C), which places TEAD1 downstream of SOX10 and NR2F2 activity.

**Figure 4.**
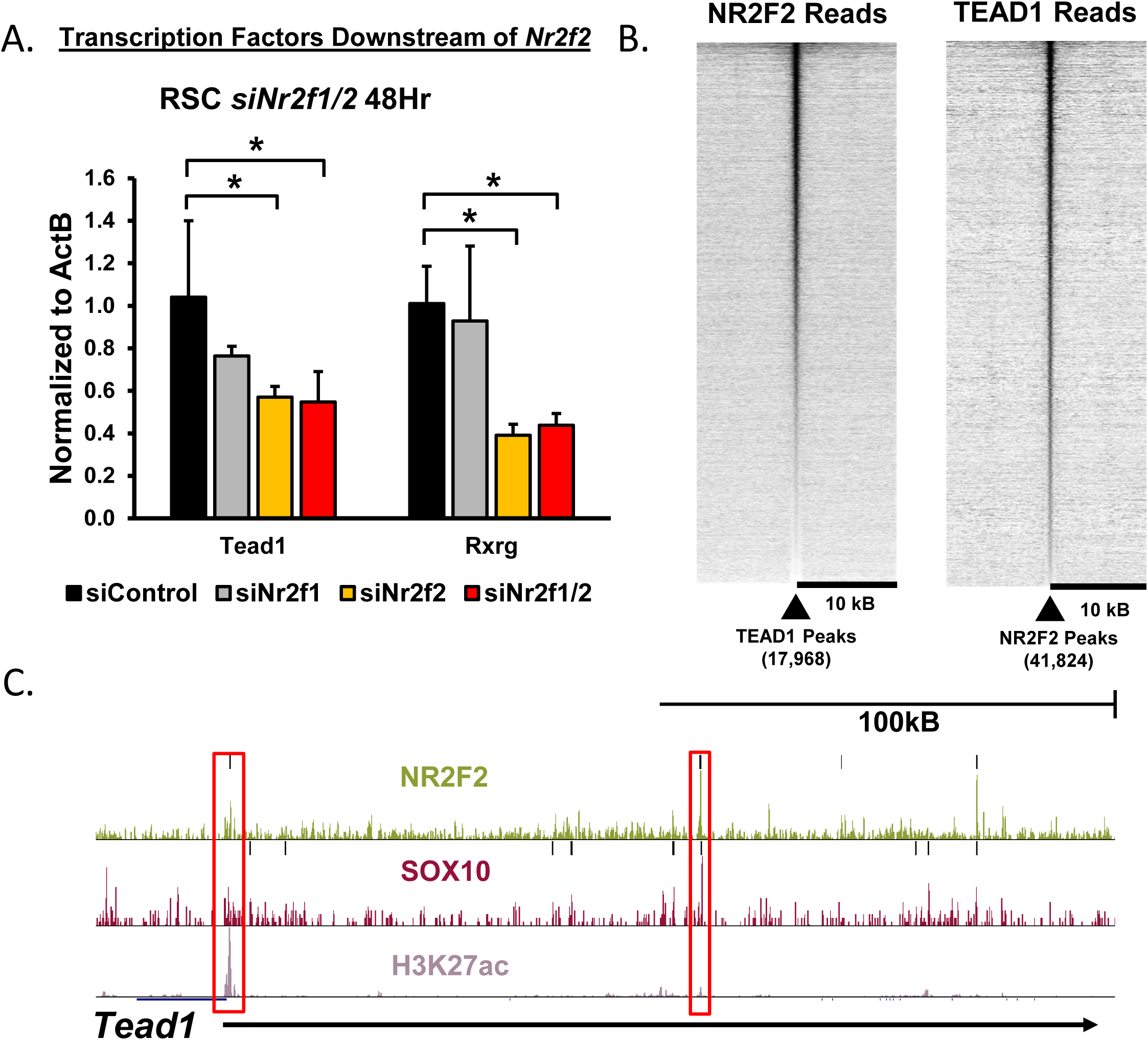
TEAD1 is Regulated by NR2F2 and Colocalizes with NR2F2. **(A)** qRT-PCR analysis of primary rat Schwann cells treated with siRNA for Nr2f1/2 (n=3, * = p<0.05) shows downregulation of transcription factor genes *Tead1* and *Rxrg*. **(B)** Heatmaps show read density of NR2F2 Cut&Run assays centered on TEAD1 peaks, and inversely the distribution of TEAD1 Cut&Run reads centered on NR2F2 peaks **(C)** Cut&Run tracks of RXRG and NR2F2 in the S16 Schwann cell line are shown, along with a ChIP-seq track of H3K27ac. Binding profiles across the *Tead1* gene reveal several enhancers bound by RXRG and NR2F2.

To determine if NR2F2 may co-regulate genes with TEAD/YAP/TAZ transcription factor, we compared genes downregulated in *siNr2f1* and *Nr2f2* with those downregulated in RNA-seq data from YAP/TAZ knockout mice (Poitelon et al., 2016), which identified a set of 38 common target genes for NR2F and TEAD/YAP/TAZ (p < 0.01, Supporting Information Table S5). This gene list includes nmSC genes such as *Scn7a*, suggesting mutual downstream gene targets between NR2F2 and TEAD1. In addition, we found that one of the gene ontology categories enriched in our NR2F target genes was the HIPPO pathway (Supporting Information Figure S2).

Although we had previously found enrichment of TEAD motifs in SOX10-bound enhancers (Lopez-Anido et al., 2015), we further hypothesized that NR2F2 cooperates with TEAD1 at nmSC enhancers. Cut&Run assays for TEAD1 in the S16 Schwann cell line revealed co-enrichment of NR2F2 reads across TEAD1 peaks, and conversely, TEAD1 reads across NR2F2 peaks (Fig. 4B). As one example, a gene enriched in nmSC is *Lama4* encoding laminin α4, which is required for proper sorting of axons and nmSC formation (Rasi et al., 2010). *Lama4* is downregulated by *Nr2f1/2* siRNA and is proximal to an enhancer that binds TEAD1, NR2F2, and SOX10 (Fig. 5B).

**Figure 5.**
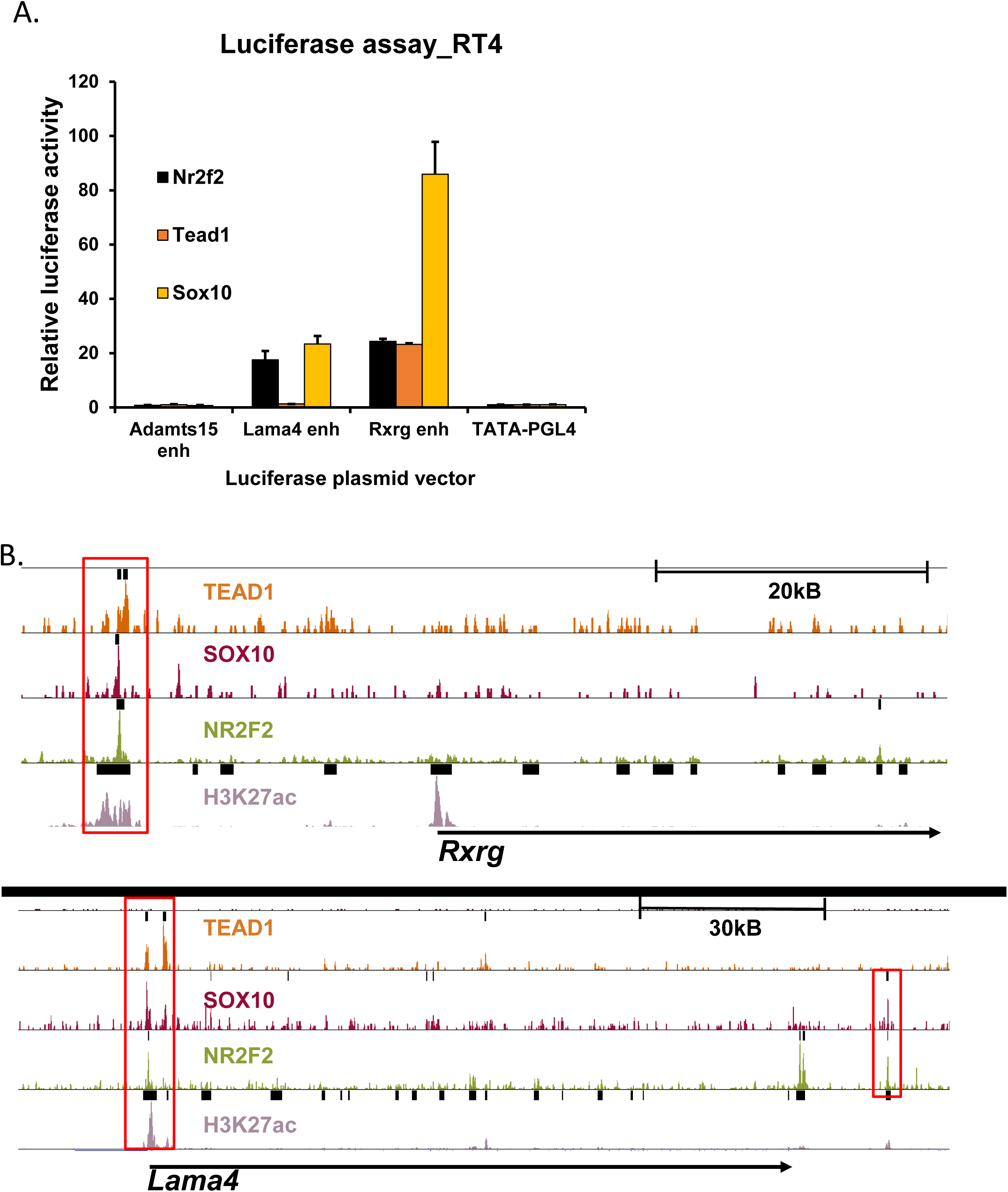
TEAD1 binding to nonmyelinating Schwann cell enhancers. **(A)** Reporter constructs of nonmyelinating Schwann cell genes were co-transfected with plasmids expressing *Sox10*, *Nr2f2*, *Rxrg*, or *Tead1*, in RT4 Schwann cells. Enhancers for *Lama4* and *Rxrg* are activated by overexpression of SOX10 and NR2F2 (n=2). **(B)** Cut&Run tracks of TEAD1, RXRG, and NR2F2 in the S16 Schwann cell line, along with a ChIP-seq tracks of SOX10 in sciatic nerve and H3K27ac in S16 cells highlight an upstream enhancer in *Rxrg*, and two upstream enhancers and one intragenic enhancer in nmSC gene, *Lama4*.

### Regulation of RXR-gamma by NR2F2 and SOX10

Although NR2F2 can form homodimers, it can also heterodimerize with other nuclear receptors in the RAR/RXR receptor family (Cooney et al., 1992; Kliewer et al., 1992; Tran et al., 1992). Intriguingly, one of the target genes reduced by siNr2f1/2 was Retinoic X receptor gamma (*Rxrg*), which is expressed in developing Schwann cells in a single cell survey of neural crest development (Kastriti et al., 2022) and is also regulated by SOX10 (Fogarty et al., 2020). We accordingly found a SOX10-bound enhancer 30 kb from the *Rxrg* transcription start site, which also had strong binding of NR2F2 (Fig. 5B). RNA-seq data from purified mSC and nmSC in post-natal day 5 mouse nerve (Gerber et al., 2021) showed that *Rxrg* is strongly expressed in nmSC but almost absent in mSC.

### NR2F2 regulates Nonmyelinating Schwann Cell Enhancers

To determine if NR2F2 and TEAD1 can activate enhancers proximal to nmSC genes, reporter constructs for enhancers proximal to nmSC genes *Rxrg*, *Adamts15*, and *Lama4*, were co-transfected with overexpression plasmids for NR2F2, TEAD1, and SOX10 in RT4 Schwann cells (Fig. 5A). SOX10 expression resulted in the strongest activation of the *Rxrg* enhancer, while also activating *Lama4* to a lesser extent. Although the Adamts15 enhancer was not responsive in this assay, NR2F2 expression activated enhancers of *Rxrg* and *Lama4*.

### A human gene regulatory network in nonmyelinating Schwann cells

In parallel to these studies, we have developed gene regulatory networks for human Schwann cells using recently published single cell RNA-seq and ATAC-seq data (Zhang et al., 2021; Avraham et al., 2022) using NetREm methodology (Khullar et al., 2025). The resulting model provides distinct transcription factor-target gene (TG) regulatory networks and transcription factor-transcription factor (TF-TF) coordination networks, which is applied for each target gene (TG) in the given Schwann cell sub-type and aggregated to reveal cell-type networks in mSCs and in nmSCs. For each TG, NetREm inputs single-cell gene expression data for candidate TFs based on motifs present within regulatory elements identified by ATAC-seq (Zhang et al., 2021). Then, NetREm utilizes a Protein-Protein Interaction (PPI) network to perform network-constrained regularization to predict TG expression in mSC and nmSC based on the gene expression levels of the TFs. This model can use the human single cell data to infer the importance of NR2F2 and collaborating TFs in mSC and nmSC.

As shown in Figure 6A, NetREm reveals a network of TF-TG regulatory links, which is used to derive SC-sub-cell-type-specific TF-TF coordination (given by coordination scores *B*) for each TG. The *B* coordination score for a TF-TF link reflects the frequency by which they co-regulate TGs in the network. Overall, TF-TG regulatory links were uncovered for 8,950 TGs (8,943 TGs have at least 2 TFs) in mSCs and 5,207 TGs (all have at least 2 TFs) in nmSCs, with 3,841 TGs shared between the networks. For each TF-TF link, the coordination scores across all TGs with at least 2 selected TFs were calculated, resulting in 22,809 non-zero undirected TF-TF coordination links 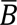 in mSCs and 24,795 such links in nmSCs. The matrix shown in Figure 6A shows the cell-type-specific 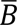 coordination score (i.e. percentile ranking) for TF-TF links involving NR2F2, RXRG, and other transcription factors important for Schwann cell development (SREBPF1, SOX10, EGR2, YY1, and TEAD1) within mSC and nmSC’s. The differential enrichment of the TF-TF links in nmSC vs mSC is shown in the network diagram (Figure 6B), which shows that the predicted human gene regulatory network for mSC genes is enriched for EGR2-SOX10 linkages, whereas the nmSC gene network is highly enriched for TF pairs involving SOX10, NR2F2, RXRG, and TEAD1.

**Figure 6.**
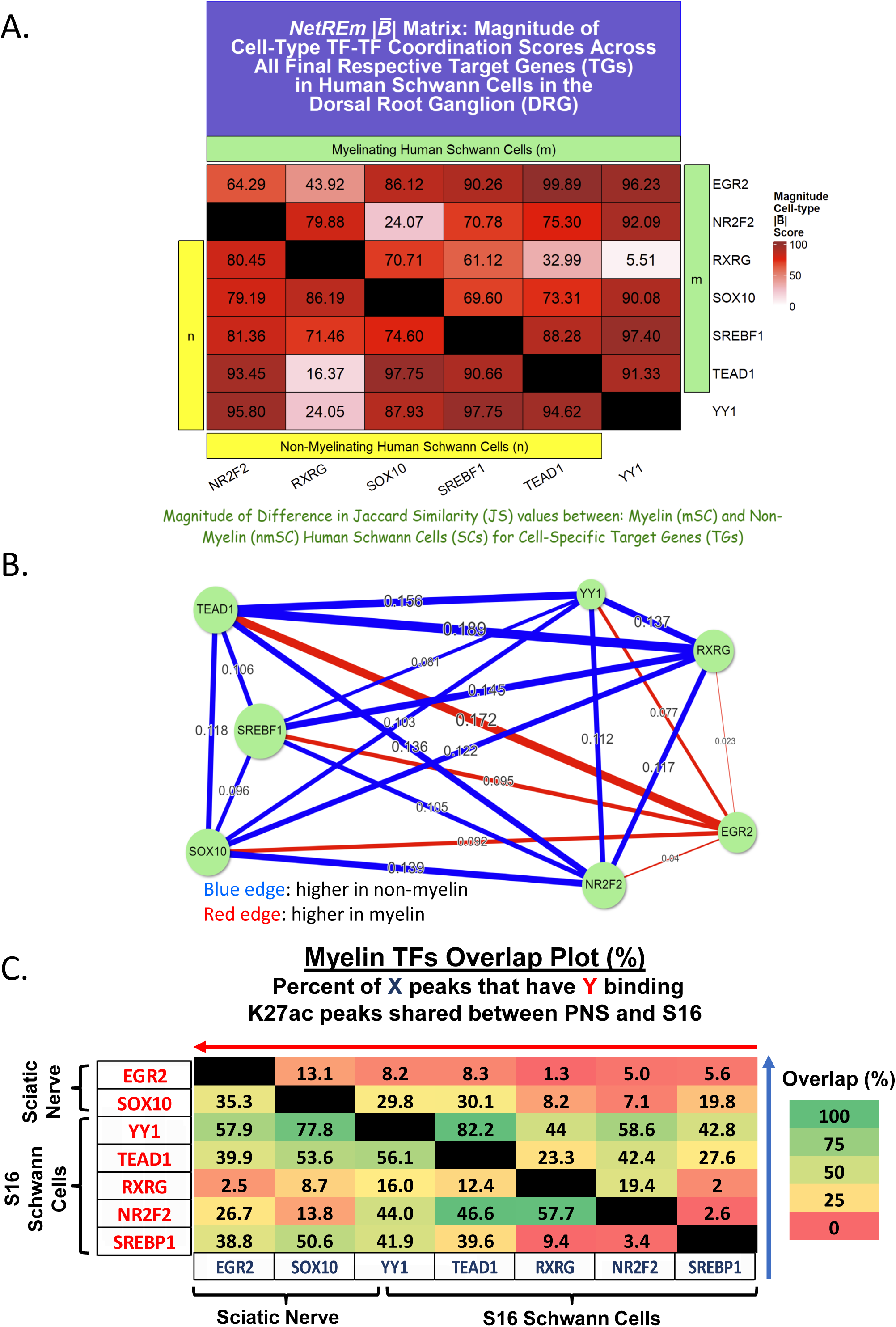
Human Transcription Factor Interactions in myelinating and nonmyelinating Schwann cells. **(A)** Using the Netrem analysis of human single cell RNA-seq and ATAC-seq data, a heatmap of relative percentile of 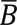 coordination scores for 7 core SC TFs (EGR2, NR2F2, RXRG, SOX10, SREBF1, TEAD1, and YY1) is displayed, with the exception of mSC-specific TF EGR2, which is not expressed in nmSCs. TEAD1 has very strong coordination (above 93.34 percentile) with these respective TFs across myelinating and nonmyelinating SCs. RXRG has stronger relative coordination links with these TFs in nmSCs than in mSCs. **(B)** TF-TF regulatory networks (based on the final NetREm model that predicts TG gene expression levels) are shown for 7 core SC TFs for TGs that are specific to each SC type: 3,611 mSC-specific TGs and 785 nmSC-specific TGs. Displayed numbers are calculated by counting the number of common TGs shared by a pair of TFs in that SC type and then compute the Jaccard Similarity (JS) of TGs between a pair of TFs. Blue lines signify higher relative co-regulation in nmSCs. The network shows high relative co-regulation between RXRG and NR2F2 in nmSCs (JS = 0.2 with 16 common nmSC-specific TGs vs. none in mSCs), which is consistent with our findings of higher 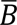 scores and relative coordination score percentiles in nmSCs compared with mSCs. In contrast, EGR2 has higher JS values with the 6 remaining SCs in mSCs (and consequently has red edges) as it is not expressed in nmSCs. **(C)** Overlap matrix of called peaks for chromatin immunoprecipitation transcription factors in mouse nerve (EGR2, SOX10, and SREBP1) and Cut&Run transcription factors in S16 rat Schwann cells (NR2F2, RXRG, TEAD1, YY1). The plot displays percentages of peaks from one factor that overlap with another factor. Cells are shaded with heatmap colors from green (more preferential overlap) to red (less preferential overlap).

Since this network was based purely on human genomic data without any inputs of our rat ChIP/Cut&Run data, this provides an independent validation of key TF-TF combinations in the nmSC network. The human network was compared with rat binding data that we have generated for several myelin transcription factors (Khullar et al., 2025), including SREBF1, which is an important regulator of lipid biosynthesis in myelin formation (Verheijen et al., 2009), as well as YY1, which is also required for Schwann cell development (He et al., 2010). By combining these data sets, we calculated the overlap of binding sites using our S16 Cut&Run data (NR2F2, RXRG, TEAD1, YY1) and in vivo ChIP-seq data (SOX10, EGR2, SREBP1), and the preferential pairings are apparent in this analysis (Figure 6C). For example, NR2F2 and RXRG have relatively low overlaps with the transcription factors important for mSC (e.g. EGR2 and SREBP1). However, the overlap is much greater for NR2F2 and RXRG with transcription factors involved in both mSC and nmSC development: SOX10 and TEAD1.

## Discussion

Although transcription factors exert a powerful influence on cellular differentiation, single transcription factors are limited in their ability to modulate cell-specific gene programs on their own. It has been well established that gene regulation emerges from combinatorial activation of enhancers and promoters by cooperating transcription factors (Arnone and Davidson, 1997; Heinz et al., 2010; Rada-Iglesias et al., 2012). Therefore, a more comprehensive understanding of cellular differentiation requires an ability to elucidate combinatorial networks of several types of transcription factors, most of which are not exclusively expressed in a specific cell type.

Mature Schwann cells can take two major forms: mSC or nmSC, though recent studies have identified more complex diversity in Schwann cell populations by identifying distinct myelinating subtypes (Yim et al., 2022). Although SOX10 is persistently expressed in Schwann cells and their precursors, there are many gaps in our understanding of cooperative transcriptional partners that drive Schwann cell precursors to mature Schwann cells compared to other neural crest lineages, and those that differentiate mSC and nmSC (Finzsch et al., 2010).

Studies of Schwann cell differentiation have identified a series of key transcription factors in addition to SOX10, such as EGR2/KROX20, POU3F1/OCT6, YAP/TAZ/TEAD1, SREBF1, YY1, and CTCF, which play crucial roles at various stages of Schwann cell differentiation (Topilko et al., 1994; Kuhlbrodt et al., 1998b; Kuhlbrodt et al., 1998a; Nagarajan et al., 2001; Jaegle et al., 2003; Jang et al., 2006; LeBlanc et al., 2007; Ryu et al., 2007; Yin et al., 2007; He et al., 2010; Lopez-Anido et al., 2016; Grove et al., 2017; Wang et al., 2020a). Accordingly, there are many examples of individual enhancers in myelin-associated genes that are cooperatively activated by several transcription factors (Denarier et al., 2005; Reiprich et al., 2010; Jones et al., 2012; Srinivasan et al., 2012; Lopez-Anido et al., 2016; Pantera et al., 2018; Sock and Wegner, 2019; Pantera et al., 2020).

The importance of combinatorial transcription factors in Schwann cell gene regulation emerges from several lines of evidence. First, knockouts of single transcription factors can decrease expression of abundantly expressed myelin genes, but target gene levels rarely decrease to the low baseline levels found in other cell types (Le et al., 2005; Verheijen et al., 2009; Poitelon et al., 2016). Second, there is no known transcription factor that is exclusively expressed in Schwann cells, as all those listed above are expressed in other cell types. Therefore, investigation of target gene relationships requires a comparably combinatorial analysis in order to fully reveal gene regulatory networks. Our analysis uses a novel approach to uncover a Schwann cell-specific transcriptional network that encompasses a large number of enhancers and highlights the role of nuclear receptors in nmSC.

We identified nuclear receptors NR2F1 and NR2F2 as prime candidates for cooperative transcriptional partners of SOX10 because they are co-expressed from neural crest to mature Schwann cells (Rada-Iglesias et al., 2012; Lopez-Anido et al., 2015; Gerber et al., 2021; Bonnamour et al., 2022). Additionally, we had found that knockdown of *Nr2f1* and *Nr2f2* transcripts downregulated some Schwann cell developmental and myelin genes (Lopez-Anido et al., 2015). Surprisingly, RNA-seq analysis of the *Nr2f1 and Nr2f2* siRNA knockdown in the S16 Schwann cell line led to downregulation of a gene set that is highly enriched in nmSC. Our data further indicate that NR2F1 and NR2F2 interacts with SOX10 to regulate nmSC genes, and its expression is highest in nmSC nuclei.

Binding partners of nuclear receptors in nmSC could also be involved in specifying between the two types of mature Schwann cells. One of the genes downregulated by *Nr2f1 and Nr2f2* siRNA is RXRG, which is a related nuclear receptor that can heterodimerize with NR2F2. Single-cell RNA-seq data revealed *Rxrg* is also preferentially expressed in nmSC (Gerber et al., 2021).

The initial identification of NR2F binding sites was based on SOX10-bound enhancers (Lopez-Anido et al., 2015), and SOX10 is required for both mSC and nmSC development (Finzsch et al., 2010). Similarly, both mSC and nmSC require the HIPPO pathway of transcriptional regulators, including co-activators Yes-Associated Protein (YAP) and Transcriptional coactivator PDZ-motif (TAZ), which interact with TEADs to regulate downstream targets (Harvey et al., 2003; Wu et al., 2003; Chen et al., 2010). In Schwann cells, expression of YAP and TAZ are required for axonal sorting and myelin development (Poitelon et al., 2016; Grove et al., 2017). A specific knockout of *Tead1* in Schwann cells causes radial sorting defects and impairs the formation of Remak bundles by nmSC (Grove et al., 2024). Consistent with our previous analysis of SOX10-bound enhancers in Schwann cells (Lopez-Anido et al., 2015), we found extensive colocalization of TEAD1 and SOX10. Moreover, NR2F2 and TEAD1 have strong colocalization as well, suggesting that interactions of NR2F2/RXRG with SOX10 and TEAD1 could be a major determinant of nmSC transcriptional regulation.

While our studies were based on genomic localization data in rat peripheral nerve in vivo and the S16 Schwann cell line, we were also able to take advantage of parallel studies of single cell data from human Schwann cells (Zhang et al., 2021; Avraham et al., 2022). An independent analysis of these data as shown in Figure 8 revealed distinct transcriptional networks for mSC and nmSC, and these networks similarly showed preferential regulation of nmSC genes by RXRG and NR2F2. The analysis also revealed extensive coordination between RXRG/NR2F2 and SOX10/TEAD1 in the nmSC network, consistent with our rat Schwann cell data.

A recent paper identified a role for NR2F2 in promoting Schwann cell differentiation *in vitro* by induction of *Egr2* (Han et al., 2023). The authors found that administration of db-cAMP increases the expression of *Nr2f2* in primary rat Schwann cells and that knockdown of *Nr2f2* in the presence of db-cAMP prevented induction of important myelin genes such as *Egr2*, *Pou3f1*, and *Mpz.* Our data did not highlight an overall role for NR2F2 in promoting the EGR2-regulated myelin gene network in the S16 cell line. In contrast, our data suggest a role in regulation of nmSC development. However, NR2F factors may associate with different transcription factors at different stages to transition regulation from myelin genes to nonmyelinating Schwann cell genes. Indeed, another potential role of NR2F factors emerged from characterization of the “Spot” mouse model that came from a forward genetics screen for neural crest defects. The Spot model mimics Waardenburg syndrome due to an insertion mutant in the *Nr2f1* gene (Bergeron et al., 2016), causing transcript overexpression and premature Schwann cell differentiation in the neural crest (Bonnamour et al., 2022). Intriguingly, NR2F1 overexpression switches melanocyte-fated cells to Schwann cells in the skin. *Nr2f1* and *Nr2f2* are co-expressed in the neural crest during development, and RNA-seq data from Spot neural crest cells enriched NR2F2 as a paralogous co-regulator of gliogenesis, with both receptors targeting Hedgehog and Notch pathway genes (Bonnamour et al., 2022). In light of these gain-of-function data, a full analysis of NR2F2 regulation will require targeted disruption of NR2F2 and perhaps NR2F1 in knockout mice.

We developed the application of Cut&Run assays (Skene and Henikoff, 2017) to probe the cooperative interactions of different transcription factors in the S16 Schwann cell line. This provided complementary data to our previous studies employing ChIP-seq, which we had used in rat peripheral nerve for transcription factors and histone modifications (Jang and Svaren, 2009; Jang et al., 2010; Hung et al., 2012; Jones et al., 2012; Srinivasan et al., 2012; Hung et al., 2015; Ma et al., 2015). Cut&Run assays have some advantages compared to ChIP-seq, since less cells are required, and the technique seems to work with a broader range of antibodies (Skene and Henikoff, 2017).

The combinatorial interactions of SOX10 with NR2F2 and other partners is presumably required to drive the Schwann cell-specific landscape of enhancers required for differentiation. While studies of specific enhancers are valuable, it has now become possible to understand how these transcriptional regulators interact with enhancers and promoters across the genome. Using Cut&Run, we have defined transcriptional clusters that regulate nmSC genes and have uncovered potential targets that can shift Schwann cells between nonmyelinating and myelinating fates.

### Experimental Procedures

#### Cut and Run (C&R)

S16 Schwann cells were cultured in three 10cm plates to yield 10 million cells. Plates were washed with PBS containing protease inhibitor cocktail, then washed with nuclear extraction buffer (20mM HEPES pH 7.9, 10mM KCl, 1.5mM MgCl2, 0.5mM Spermidine, 10% Glycerol, 0.3% Triton X-100) to isolate nuclei. Cell nuclei were bound to Concanavalin A beads using the Cut&Run assay kit (Cell Signaling Technology #86652). Nuclei were permeabilized with digitonin, then incubated with protein AG-micrococcal nuclease (PAG MNase). The following antibodies were used: NR2F2 Perseus Proteomics PP-H7147-00), RXRG (Developmental Studies Hybridoma Bank PRCP-RXRG-5H4), and TEAD1 (BD Biosciences 610923). After removal of Concanavalin-bound nuclei, DNA was purified from the remaining solution using the Cell Signaling DNA purification kit (Cat# 14209S). Cut&Run samples were processed using the Illumina NovaSeq6000 system and NEBNext Cut&Run library prep. Cut&Run data have been deposited in NCBI GEO under accession number: GSE247955.

#### Cell Culture and siRNA

S16 Schwann cells and Primary Rat Schwann cells (RSCs) were cultured in 6-well plates in Corning DMEM containing 4.5 g/L glucose and L-glutamine without sodium pyruvate (#10-017-CV) and 5% bovine growth serum (Hyclone). S16s were transfected with control siRNA (IDT DS NC1), or siRNAs for *Nr2f1* (IDT, RNC.RNAI.N031130.12.3), *Nr2f2* (IDT RNC.RNAI.N080778.12.1), using Mirus Transit X2 (MIR 6000), as described (Lopez-Anido et al., 2015).

A repeat of the Nr2f experiment was done with independent Nr2f siRNA’s (n=3) from IDT using the AMAXA 4D nucleofector with the AMAXA SE Cell Line 4D Nucleofector Kit L (Lonza). *Nr2f1* siRNA: mm.Ri.Nr2f1.13.1

5’-CUUCAAGAAUUAAAAUUGAAGUGAA-3’;

5’-AAGAAGUUCUUAAUUUUAACUUCACUU-3’

*Nr2f2* siRNA: mm.Ri.Nr2f2.13.1

5’-GAUGUUACAAGUUUGCUAAAAGAAG-3’

5’-UCCUACAAUGUUCAAACGAUUUUCUUC-3’

S16 Schwann cells were transfected with siRNAs for control (IDT DS NC-1), *Nr2f1* (mm.Ri.Nr2f1.13.1), *Nr2f2* (mm.Ri.Nr2f2.13.1), and *Sox10*. This was done using nucleofection (AMAXA 4D nucleofector with the AMAXA SE Cell Line 4D Nucleofector Kit L (Lonza)). There were four experimental groups: control, si*Nr2f1/Nr2f2*, si*Sox10*, and si*Nr2f1/Nr2f2/Sox10*. Each group had the same amount of siRNA. The control group had 9.9ul of NC-1 siRNA. si*Nr2f1/Nr2f2* had 3.3ul of *Nr2f1* siRNA, 3.3ul of *Nr2f2* siRNA, and 3.3 of NC-1 siRNA. si*Sox10* had 3.3ul of *Sox10* siRNA and 6.6 ul of NC-1 siRNA. The triple knockdown group had 3.3ul of each siRNA (*Nr2f1*, *Nr2f2*, and *Sox10*). Forty-eight hours later, the RNA was extracted using Trizol reagent (Invitrogen) and RNA Clean & Concentrator kit (Zymo Research).

*Sox10* siRNA: s131239 (Applied Biosystems)

5’-AGGUCAAGAAGGAACAGCAtt-3’

5’-UGCUGUUCCUUCUUGACCUtg-3’

RT4-D6P2T (RT4) rat Schwann cells were plated in 6-well dishes at a density of ∼2×10^5 per well a day before transfection. 250ng of Nr2f2 plasmid DNA was transfected into cells using the TransIT-X2 (Mirus, MIR 6000), following the manufacturer’s protocol. An empty vector was used as a negative control. Forty-eight hours later, the RNA was extracted using Trizol reagent (Invitrogen) and RNA Clean & Concentrator kit (Zymo Research).

#### RNA Extraction, Reverse transcription, and Quantitative PCR

6-well plates of Schwann cells were homogenized in 0.5 ml of Trizol (Thermo Fisher, #15596026) per well for 5 minutes on a rotator at low speed. Lysate was placed in RNAse free microcentrifuge tubes, 0.2 ml of chloroform was added, and tubes were centrifuged at 12,000g for 15 minutes at 4°C. The aqueous layer was purified using the Zymo RNA Clean & Concentrator kit protocol (catalog #R1014, Zymo Research) at step 2. RNA samples were analyzed using quantitative RT-PCR with the indicated primer sets (Table 1) using SYBR Green Master Mix (Applied Biosystems) on the StepOnePlus system (Applied Biosystems). All transcripts were normalized to *Actb* or *Yy1* and respective control siRNAs and plasmids using the ΔΔCT method.

1000 ng total RNA was sent to Novogene for library preparation after PolyA selection and Illumina sequencing (Novaseq 6000). Messenger RNA was purified from total RNA using poly-T oligo-attached magnetic beads. After fragmentation, the first strand cDNA was synthesized using random hexamer primers, followed by the second strand cDNA synthesis.

Reads were aligned to the Rn6 genome using Hisat2 (Mortazavi et al., 2008). FeatureCounts v1.5.0-p3 was used to count the reads numbers mapped to each gene. And then FPKM of each gene was calculated based on the length of the gene and reads count mapped to this gene. FPKM, expected number of Fragments Per Kilobase of transcript sequence per Millions base pairs sequenced. Data were analyzed using DESeq2 v1.20.0 (Anders et al., 2013), and the resulting p-values were adjusted using the Benjamini and Hochberg’s approach for controlling the false discovery rate. Genes with an adjusted P-value <=0.05 found by DESeq2 were assigned as differentially expressed. RNA-seq data are deposited in NCBI GEO under accession number GSE252428. The 2^nd^ repeat of the siNr2f RNA-seq experiment was sequenced at the UW Biotechnology Center. RNA sequencing results were aligned to the Rn6 genome using STAR 2.7.11b (Dobin et al., 2013). Data in the form of raw read counts were analyzed using DESeq2 in R version 2023.12.1+402.

Gene lists for myelinating and nonmyelinating Schwann cells were extracted from the Sciatic Nerve Atlas (Gerber et al., 2021). Replicates of myelinating and nonmyelinating Schwann cell samples (n=4) were averaged, then sorted for an RPKM value greater than 5 (Supporting Information Table S4). The lists were then sorted for nonmyelinating Schwann cell genes over myelinating Schwann cell gene with a 2-fold or greater differential cutoff value. P-values for gene list comparisons were generated using the Fisher’s exact test (Supporting Information Tables S3 and S5).

#### Luciferase Assay

The following Schwann cell enhancers, designated by mouse mm10 coordinates were cloned into a pGL4 luciferase plasmid containing a minimal promoter: Adamts15 chr9:30,896,450-30,897,040; Lama4 chr10:38,553,551-38,554,030; Rxrg: chr1:167,566,364-167,566,801; RT4-D6P2T (RT4) rat Schwann cells were plated in 6-well dishes at a density of ∼2×10^5^ per well a day before transfection. RT4 Schwann cells were co-transfected with the gene overexpression plasmids, the promoter-containing reporter vectors, and a pNL1.1 vector containing TK promoter using TransIT-X2 transfection reagent (Mirus, #MIR6004).

After a 48-hour transfection period, the cells were harvested using the Passive Lysis buffer (Promega, #E1910). Subsequently, a Nano-Glo Dual-Luciferase assay was performed using the following steps. To perform the Luciferase assay, the Luciferase assay buffer (84mM Tris-MES (pH 7.8), 20mM Magnesium Acetate, and 4mM ATP) and 1mM D-Luciferin solution were added to the cell lysates in a 1.5ml tube. The Firefly luciferase activity was measured using the GloMax 20/20 instrument (Promega). The Nano-Glo luciferase assay was performed using the Nano-Glo assay kit (Promega, N1110). Firefly luciferase activity was then normalized to the Nano-Glo activity.

### Immunohistochemistry

Freshly dissected 4-month old rat sciatic nerve was placed in O.C.T. (Tissue-TeK Cat# 4583) and snap-frozen in liquid nitrogen. 10uM sections were dried for 5min, fixed with 4%PFA in PBS, then blocked (5% donkey serum, 1% BSA, 0.5% TritonX 100 in PBS) for 1 hour. Sections were washed in PBS for 4min twice, then incubated with a primary antibody solution (5% donkey serum, 1% BSA, 0.1% TritonX 100 in PBS) overnight at 4°C. Primary antibodies used were Perseus Proteomics NR2F2 (PP-H7147-00, 1:500) and Millipore NCAM1 (AB5032, 1:500). Samples were washed in 0.1% Tween20 in PBS for 5min twice, then washed with secondary antibodies diluted in PBS for 1 hour at room temperature. Secondary antibodies used were Invitrogen AlexaFluor 488 Donkey anti-Rabbit (Cat# A21206) and Invitrogen AlexaFluor 546 Donkey anti-Mouse (Cat#A10036). Slides were washed again with 0.1% Tween20 for 5min twice. After excess moisture was removed, slides were mounted with Vectashield Plus Antifade with DAPI (Cat# H-2000). Samples were imaged using the Leica Stellaris 8 confocal with the 20x air and 63x oil objectives. Post-processing was performed using Leica Lightning Process for deconvolution and thresholding for segmentation.

#### Bioinformatics

Cut&Run FastQ files were mapped to reference genome rn5 using Bowtie2 (Langmead et al., 2009; Langmead and Salzberg, 2012) to produce Binary Alignment Map (BAM) files. BAM files were filtered for mapped reads using BamTools (Quinlan and Hall, 2010; Barnett et al., 2011) and sorted into called peaks using MACS2 (Zhang et al., 2008; Feng et al., 2012; Liu, 2014). BedTools bamCoverage generated bedgraphs of ChIP-seq samples. Heat maps were created via EAseq (Lerdrup et al., 2016). Data processing was performed in a cloud-based manner through GalaxyBiostars (Afgan, 2018). ChIP-seq tracks were visualized using the University of California, Santa Cruz Genome Browser (Kent et al., 2002). Previous ChIP-seq datasets for H3K27ac (Hung et al., 2015) and SOX10 (Lopez-Anido et al., 2015) are available at GEO using the following accession numbers: GSE63103 and GSE64703. MACS2 called peaks for NR2F2 and RXRG were processed through HOMER to identify enriched motifs (Heinz et al., 2010). For YY1, RXRG, NR2F2, TEAD1, and H3K27ac the Cut&Run (Cleavage Under Targets & Release Using Nuclease) assays in the rat S16 Schwann cell (SC) line and the SREBP1 ChIP-seq data in sciatic nerve are available from GEO (GSE247955).

RNA-seq heatmaps were generated from siNr2f1/2 RNA-seq data (GEO: GSE252428) using R (Team, 2023) and the pheatmap package (Kolde). RPKMs per sample were log2 transformed and sorted by row means.

For regulatory network prediction in human nmSC and mSC, we use NetREm (Khullar et al., 2025), which leverages multimodal data to predict functionally-coordinating TFs for target gene (TG) co-regulation. NetREm incorporates published scATAC-seq data (Zhang et al., 2021; Avraham et al., 2022), which examines adult SCs, along with human databases of Position Weight Matrices (PWMs) for TFs, TF-TF Molecular Function similarity (Zhao and Wang, 2018), and TF-TF Protein-Protein Interaction networks (PPINs) featuring direct and/or indirect PPIs (Szklarczyk et al., 2023). NetREm initially predicts candidate GRNs for each cell type by establishing TF-regulatory element-TG links based on prior knowledge of candidate TFs for each TG.

Using scRNA-seq data from human DRG by (Zhang et al., 2021; Avraham et al., 2022), NetREm applies a network regularized regression method for each TG. It selects TFs whose expression levels predict TG expression levels, subject to constraints from an input PPIN among TF predictors. For each TG in a SC cell-type, NetREm identifies final TFs that coordinate to co-regulate it, generating undirected networks of TG-specific SC cell-type TF-TF coordination, characterized by coordination scores *B* ranging from −100 to 100. A *TF*_*i*_-*TF*_*j*_ link for a given TG has *B* ≠ 0 score only if both TFs are selected as final TFs for the TG. *B*_*ij*_ > 0 implies that both TFs may cooperate (i.e. both co-repressors or co-activators), and *B*_*ij*_ < 0 suggests potential activator-repressor antagonism to regulate TG. NetREm also predicts a directed network of TF-regulatory element-TG links based on model coefficients *c*^∗^ (>0: activator, <0: repressor).

Ultimately, NetREm is applied for all TGs in the given SC sub-type to yield cell-type complementary GRNs (TF-RE-TG links for all TGs) for 8,950 TGs in mSCs and 5,207 TGs in nmSCs, with 3,841 TGs shared between the cell types. NetREm also predicts undirected cell-type TF-TF coordination network with scores 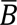 between −100 and 100 in mSCs and nmSCs.

Here 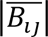 is a signed percentile, which ranks the strength of each *TF*_*i*_-*TF*_*j*_ cell-type coordination interaction relative to other TF-TF interactions. 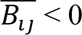 suggests activator-repressor antagonism, while a positive 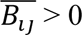 indicates potential cooperativity for TGs. Overall, NetREm identifies 22,809 TF-TF links with non-zero coordination scores (i.e. 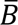 ≠ 0) for 221 final TFs in mSCs and 24,795 non-zero TF-TF links for 228 final TFs in nmSCs.

We find 5,109 mSC-specific TGs and 1,366 nmSCs-specific TGs. We calculate the Jaccard similarity (JS) index for TF pairs based on the SC-subtype-specific TGs as follows:

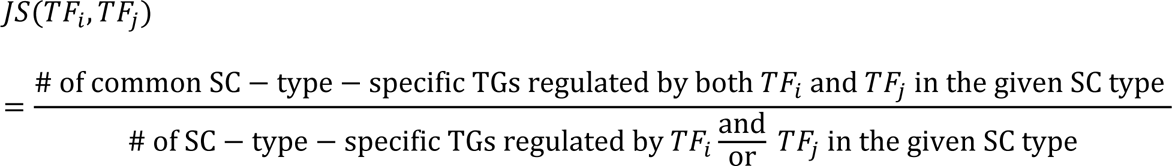

## Data Availability

Cut&Run data have been deposited in NCBI GEO under accession number: GSE247955. RNA-seq data are deposited in NCBI GEO under accession number GSE252428.

## Supporting Information

This article contains supporting information.

## Acknowledgments

The authors thank the University of Wisconsin Biotechnology Center DNA Sequencing Facility for their sequencing services. We also thank Jerome Choi and Karla Knobel for technical assistance.

## Funding

This work was supported by R21 NS127432 provided by NINDS, and a core grant to the Waisman Center from NICHD (P50 HD105353).

## Supporting Information

Table S1: RNA-seq analysis of S16 cells treated with Nr2f1/Nr2f2 siRNA Deseq2 output.

Table S2: RNA-seq analysis of S16 cells treated with independent Nr2f1/2 siRNA, Deseq2 output.

Table S3: Overlap of NR2F1/2 target genes with SOX10 target genes.

Table S4: Overlap of NR2F1/2 target genes with nmSC genes.

Table S5: Overlap of NR2F1/2 target genes with YAP/TAZ regulated genes.

Figure S1. Hierarchical Clustering of Non-Myelinating Schwann Cell Genes Downregulated in by Nr2f1/2 siRNA.

Figure S2. Antibody Validation and Gene Ontology Enrichment for the HIPPO Pathway in Genes Downregulated by Nr2f1/2.

Figure S3 Overlap of non-myelinating SC genes and genes downregulated by Nr2f1/2 siRNA in S16 cells.

## Notes

https://www.ncbi.nlm.nih.gov/geo/query/acc.cgi?acc=GSE252428

https://www.ncbi.nlm.nih.gov/geo/query/acc.cgi?acc=GSE247955

### Competing Interest Statement

The authors have declared no competing interest.

