## Supporting Information Figure for "Integration of Nuclear Receptors into a Schwann cell Gene Regulatory Network"

### Figure S1

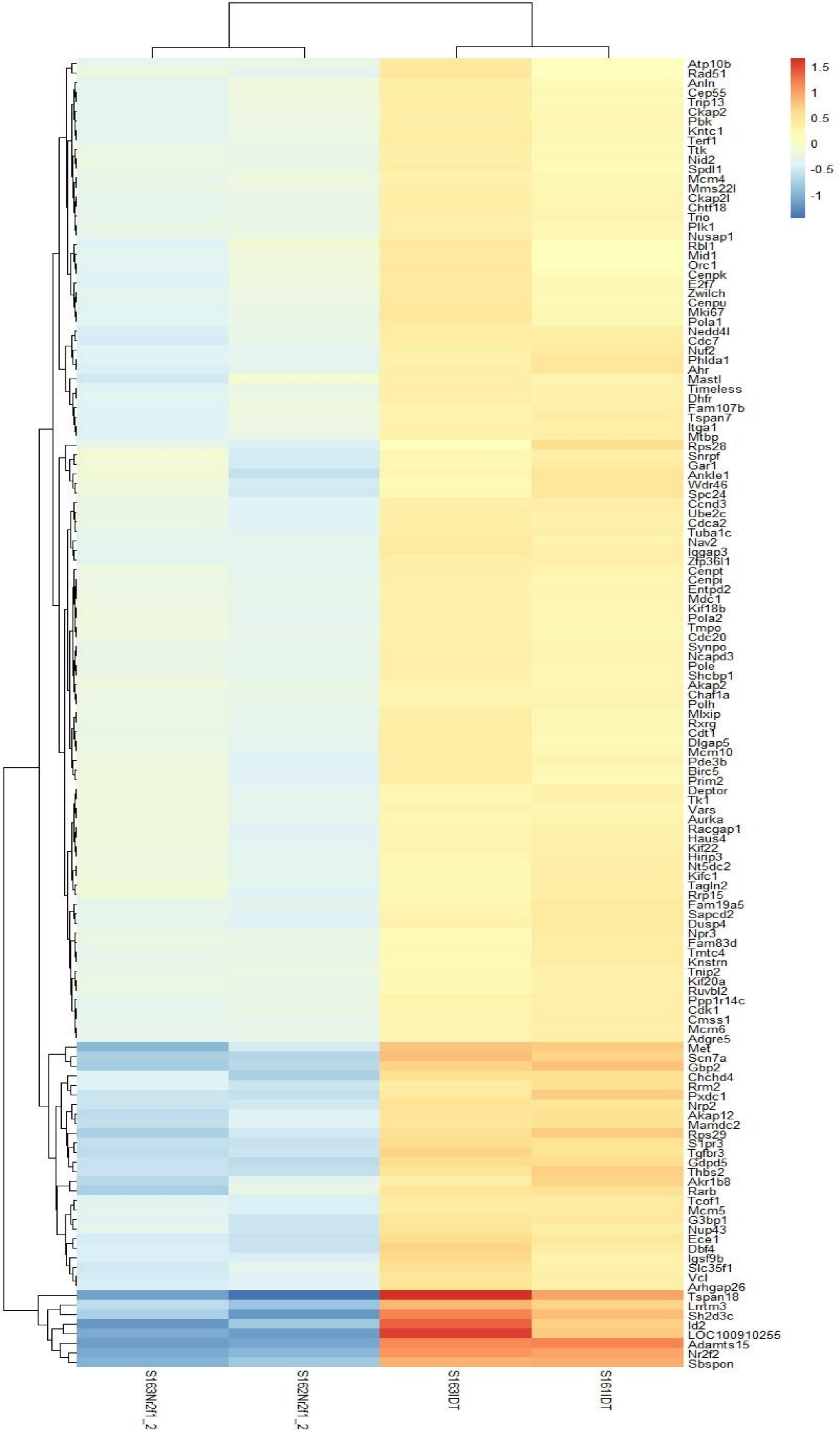


**Figure S1. Hierarchical Clustering of Non-Myelinating Schwann Cell Genes by *Nr2f1/2 siRNA.*** RNA-seq heatmap of all downregulated genes in the overlap between non-myelinating Schwann cell genes from the Suter dataset compared to the siNr2f1/2 data. Heatmap shows FPKM values log2 normalized, sorted by row means, and clustered by similar differential expression patterns.

### Figure S2

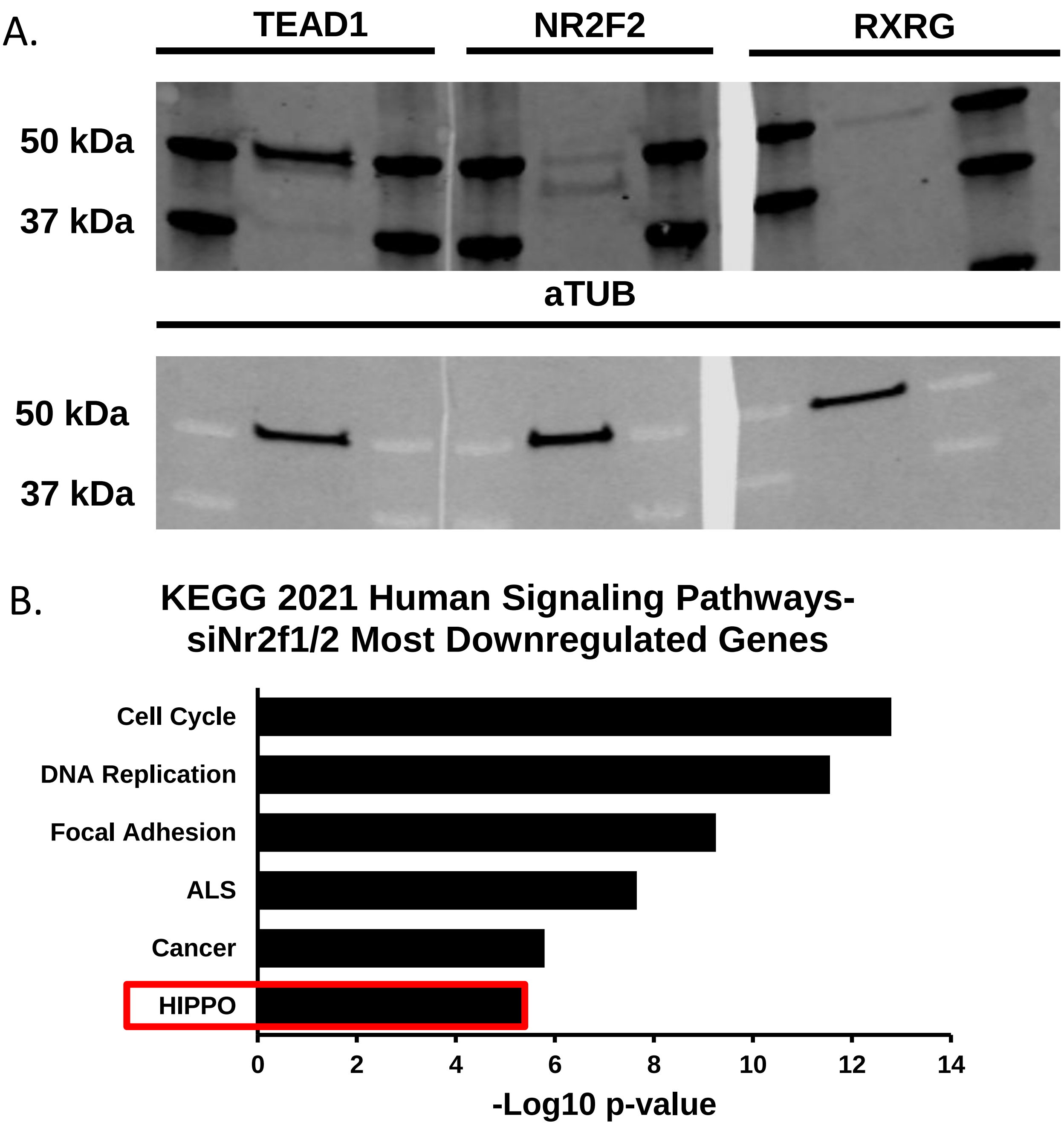


**Figure S2. Antibody Validation and Gene Ontology Enrichment for the HIPPO Pathway in Genes Downregulated by *Nr2f1/2.***
**(A)** Western blot shows protein levels of TEAD1, NR2F2, and RXRG in S16 Schwann cells.
**(B)** Gene ontology of most significantly downregulated genes in *siNr2f1/2* treated S16 Schwann cells reveals the HIPPO pathway as a target for NR2F1/2 regulation.

### Figure S3

A.

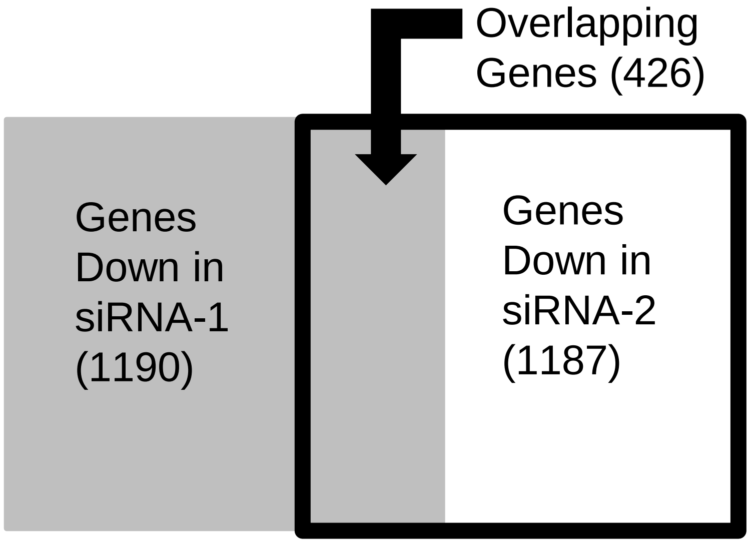


| Gene | siRNA-1 (log2FC) | siRNA-2 (log2FC) |
|---|---|---|
| Adamts15 | -2.3 | -2.3 |
| Tspan18 | -2.7 | -2.5 |
| Gbp2 | -1.5 | -1.2 |
| Tgfbr3 | -1.2 | -1.0 |
| Rxrg | -0.5 | -0.9 |

B.

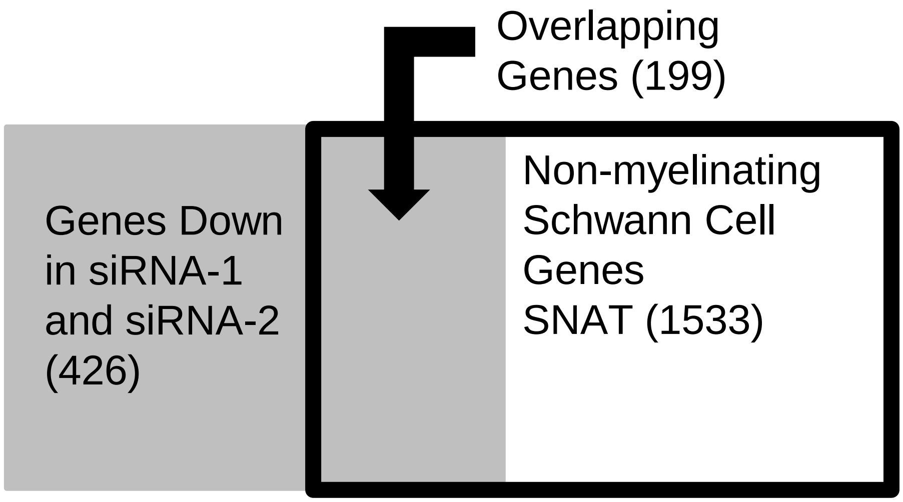


**Figure S3. Overlap of non-myelinating SC genes and genes downregulated by Nr2f1/2 siRNA in S16 cells.**
**(A)** Out of 1,187 genes knocked down (padj<0.05 and negative log2FC) using independent siRNA's for Nr2f1/2 (siRNA-2, Supporting Information Table 2), 426 overlap with the 1,190 genes reduced (padj <0.05, negative log2FC) down byoriginal siRNA for Nr2f1/2 (siRNA-1). p < 1.5e-322. **(B)** Of the 426 downregulated genes overlapping between siRNA-1 and siRNA-2, 199 overlap with 1,532 non-myelinating Schwann cell genes. p < 5e-148.
